# Chemicogenetic recruitment of specific interneurons suppresses seizure activity

**DOI:** 10.1101/291179

**Authors:** Alexandru Călin, Mihai Stancu, Ana-Maria Zagrean, John G. Jefferys, Andrei S. Ilie, Colin J. Akerman

## Abstract

Enhancing the brain’s endogenous inhibitory mechanisms represents an important strategy for suppressing epileptic discharges. Indeed, drugs that boost synaptic inhibition can disrupt epileptic seizure activity, although these drugs generate complex effects due to the broad nature of their action. Recently developed chemicogenetic techniques provide the opportunity to pharmacologically enhance endogenous inhibitory mechanisms in a more selective manner. Here we use chemicogenetics to assess the anti-epileptic potential of enhancing the synaptic output from three major interneuron populations in the hippocampus: parvalbumin (PV), somatostatin (SST) and vasoactive intestinal peptide (VIP) expressing interneurons. Targeted pre- and post-synaptic whole cell recordings in an *in vitro* hippocampal mouse model revealed that all three interneuron types increase their firing rate and synaptic output following chemicogenetic activation. However, the interneuron populations exhibited different anti-epileptic effects. Recruiting VIP interneurons resulted in a mixture of pro-epileptic and anti-epileptic effects. In contrast, recruiting SST or PV interneurons produced robust suppression of epileptiform activity. PV interneurons exhibited the strongest effect per cell, eliciting at least a five-fold greater reduction in epileptiform activity than the other cell types. Consistent with this, we found that chemicogenetic recruitment of PV interneurons was effective in an *in vivo* mouse model of hippocampal seizures. Following efficient delivery of the chemicogenetic tool, pharmacological enhancement of the PV interneuron population suppressed a range of seizure-related behaviours and prevented generalized seizures. Our findings therefore support the idea that selective chemicogenetic enhancement of synaptic inhibitory pathways offers potential as an anti-epileptic strategy.

**Significance statement:** Drugs that enhance synaptic inhibition can be effective anticonvulsants but often cause complex effects due to their widespread action. Here we examined the anti-epileptic potential of recently developed chemicogenetic techniques, which offer a way to selectively enhance the synaptic output of distinct types of inhibitory neurons. A combination of in vitro and in vivo experimental models were used to investigate seizure activity in the mouse hippocampus. We find that chemicogenetically recruiting the parvalbumin-expressing population of inhibitory neurons produces the strongest anti-epileptic effect per cell, and that recruiting this cell population can suppress a range of epileptic behaviours in vivo. The data therefore support the idea that targeted chemicogenetic enhancement of synaptic inhibition offers promise for developing new treatments.

## Introduction

The intrinsic inhibitory system of the brain is an important target for anti-epileptic drugs, since abnormalities in GABAergic function often result in epilepsy (Treiman, 2001) and drugs that enhance GABA-mediated inhibition can be potent anticonvulsants (Czapiński et al., 2005). However, because of their system-wide actions, these drugs exhibit multiple deleterious side-effects (Snodgrass, 1992; Mula, 2011). Whilst GABAergic signalling can become altered in cells within the epileptic focus (Cohen et al., 2002; Huberfeld et al., 2007; Pallud et al., 2014), inhibitory mechanisms remain effective within the ‘penumbra’ surrounding the epileptic focus and are able to oppose seizure spread (Trevelyan et al., 2006, 2007; Schevon et al., 2012; Cammarota et al., 2013). Selectively enhancing these endogenous inhibitory mechanisms therefore offers the potential to disrupt the propagation of epileptic discharges.

The region of the brain that often contains the epileptic focus in patients – the hippocampus – includes multiple subtypes of GABA-releasing interneurons, which are thought to vary in terms of their inhibitory capacity (Klausberger et al., 2003; Klausberger and Somogyi, 2008). For instance, because of their intrinsic properties and the fact that they target the perisomatic compartment of multiple pyramidal neurons, parvalbumin-expressing (PV) interneurons have been considered particularly effective at inhibiting principal neurons (Cobb et al., 1995; Freund and Buzsáki, 1996; Miles et al., 1996) and at restricting the propagation of network activity (Trevelyan et al., 2006; Cammarota et al., 2013). Meanwhile, because of their post-synaptic targeting, somatostatin-expressing (SST) interneurons have been associated with the regulation of dendritic excitability (Miles et al., 1996; Paz and Huguenard, 2015), which can then affect the spiking output of principal neurons (Lovett-Barron et al., 2012). Other interneuron subtypes, such as vasoactive intestinal polypeptide-expressing (VIP) interneurons, can mediate disinhibitory effects as well as inhibitory effects, virtue of the fact that many of their postsynaptic targets are interneurons (Acsády et al., 1996; Dávid et al., 2007; Pfeffer et al., 2013).

Consistent with these ideas, studies using optogenetic strategies to increase interneuron activity have reported promising results in terms of reducing seizure activity (Krook-Magnuson et al., 2013, 2014; Ledri et al., 2014), as well as supporting the conclusion that interneuron subtypes can exert differential effects upon seizure generation and progression (Cammarota et al., 2013; Krook-Magnuson et al., 2013; Sessolo et al., 2015; Khoshkhoo et al., 2017). The temporally-synchronous nature of optical activation can also generate counterintuitive effects, as the simultaneous recruitment of interneurons can enhance network synchronization and actually initiate epileptiform activity (Sessolo et al., 2015; Yekhlef et al., 2015; Chang et al., 2018).

A novel alternative strategy is afforded by chemicogenetic tools such as Designer Receptors Exclusively Activated by Designer Drugs (DREADDs), which use pharmacological agents to enhance or inhibit the activity of defined cell populations (Alexander et al., 2009; Roth, 2016; Smith et al., 2016). DREADDs are mutated human muscarinic receptors that can be expressed under the control of a cell-specific promoter and are not activated by endogenous ligands, but are activated by drugs such as clozapine N-oxide (CNO). Activating excitatory DREADDs, such as the human type-3 muscarinic designer receptor coupled with the G_q_ protein (hM_3_D_q_ receptor), is thought to enhance neuronal excitability by downregulating ion channels that hyperpolarise the membrane (Alexander et al., 2009).

Excitatory hM_3_D_q_ DREADDs have been used to activate interneurons in several brain regions (Hamm and Yuste, 2016; Chen et al., 2017; Funk et al., 2017; Wang et al., 2017). However, it remains unclear to what extent different subtypes of GABAergic interneurons can be modulated via DREADDs, and whether chemicogenetic control of different interneurons is a viable strategy for reducing epileptic activity. Here we demonstrate that three of the major interneuron subtypes in the hippocampus (PV-, SST- and VIP-expressing interneurons) can be successfully recruited with excitatory DREADDs. The subtypes differ however, in their capacity to increase post-synaptic inhibition in principal neurons and in their ability to reduce epileptiform network activity. Our study suggests that chemicogenetically enhancing specific interneuron populations may offer an effective anticonvulsant strategy.

## Materials and Methods

### Preparation and viral transduction of organotypic hippocampal brain slices

All animal work relating to in vitro preparations was carried out in accordance with the Animals (Scientific Procedures) Act, 1986 (UK) and under project and personal licenses approved by the Home Office (UK). Mouse organotypic hippocampal brain slice cultures were prepared from 5-to-7-day-old heterozygous or homozygous, male or female PV-Cre mice (B6;129P2-Pvalb^tm1(cre)Arbr^/J, The Jackson Laboratory), SST-IRES-Cre mice (Sst^tm2.1(cre)Zjh^/J, The Jackson Laboratory), or VIP-IRES-Cre mice (Vip^tm1(cre)Zjh^/J, The Jackson Laboratory), as described by Stoppini et al., 1991. All reagents were purchased from Sigma-Aldrich, unless stated otherwise. The brains were extracted and transferred into cold (4°C) dissection media containing Earle’s Balanced Salt Solution + CaCl_2_ + MgSO_4_ (Thermo Fisher Scientific), supplemented with 25.5 mM HEPES, 36.5 mM D-glucose and 5 mM NaOH. The hemispheres were separated, and the individual hippocampi were dissected and immediately sectioned into 400-μm-thick slices on a McIlwain tissue chopper (Mickle, UK). Slices were then rinsed in cooled dissection media, placed in 6-well plates onto sterile, porous Millicell-CM membranes, and maintained for 2-to-8 weeks in culture media containing 78.8% (vol/vol) Minimum Essential Media + GlutaMAX-I (Thermo Fisher Scientific), 20% (vol/vol) heat-inactivated horse serum (Thermo Fisher Scientific), 1% (vol/vol) B27 (Thermo Fisher Scientific), 30 mM HEPES, 26 mM D-Glucose, 5.8 mM NaHCO_3_, 1 mM CaCl_2_, 2 mM MgSO_4_⋅7H_2_O, and incubated at 35.5-to-36°C in a 5% CO_2_ humidified incubator.

After 3-to-5 days in culture, organotypic hippocampal slices were transduced with adeno-associated virus (AAV, serotype 8) containing loxP-flanked, inverted DNA sequences under the control of the human Synapsin 1 promoter (University of North Carolina Gene Therapy Center Vector Core and Addgene, USA). Viral DNA contained the double-floxed sequence for hM_3_D_q_-mCherry (Addgene #44361), which was used to target the excitatory DREADDs to specific cre-expressing populations. In control experiments and to determine interneuron expression profiles, viral DNA contained the double-floxed sequence for EGFP (Addgene #50457). Transduction was achieved by injecting viral particles (mixed with 1% wt/vol fast-green for visualization) into 5-to-10 locations along the pyramidal cell layer of the hippocampal slices. Injection pipettes were pulled from glass capillaries (1.2 mm outer diameter, 0.69 mm inner diameter; Warner Instruments) using a horizontal puller (Sutter P-97), mounted on a manual manipulator (Narishige, Japan) and monitored under a microscope (Leica S6E) coupled with an external fibre optic light source (Photonic Leica CLS 100X). A Picospritzer II system (General Valve) delivered controlled pressure pulses (5-to-10 psi for 1 s) to facilitate gradual diffusion of the viral solution into the tissue. Typical titers were ∼10^12^ IU/ml and injection volumes were ∼250 nL per slice. Feeding media was supplemented with 1% (vol/vol) antibiotic and antimycotic solution (with 10,000 units penicillin, 10 mg streptomycin and 25 μg amphotericin B per mL) for up to two feeding sessions after injection and slices were allowed at least two weeks for expression before being used.

### Electrophysiological recordings in vitro

The organotypic hippocampal slices were transferred to a recording chamber, where they were maintained at 28°C and continuously superfused with artificial cerebrospinal fluid (aCSF) containing (in mM): NaCl (120), KCl (3), MgCl_2_ (0.5-to-1.5), CaCl_2_ (2-to-3), NaH_2_PO_4_ (1.2), NaHCO_3_ (23), D-Glucose (11) and ascorbic acid (0.2). Osmolarity was adjusted to 290 mOsm and pH was adjusted to 7.36 with NaOH. Oxygen and pH levels were stabilised by bubbling the aCSF with 95% O_2_ and 5% CO_2_. Neurons within the hippocampal formation were visualised with 10x and 60x water-immersion microscope objectives (Olympus BX51WI) and targeted for single or dual-patch whole-cell recordings. Patch pipettes of 4-to-9 MΩ tip resistance were pulled from filamental borosilicate glass capillaries with an outer diameter of 1.2 mm and an inner diameter of 0.69 mm (Warner Instruments), using a horizontal puller (Sutter P-97). For current clamp recordings, pipettes were filled with a potassium-gluconate internal solution (134 mM K-gluconate, 2 mM NaCl, 10 mM HEPES, 2 mM Na_2_ATP, 0.3 mM NaGTP, 2 mM MgATP), which had been set to a pH of 7.36 using KOH, and an osmolarity of 290 mOsm. For recording post-synaptic inhibitory currents in voltage clamp, pipettes were filled with a caesium-gluconate internal solution (120 mM Cs-gluconate, 4 mM NaCl, 40 mM HEPES, 2 mM MgATP, 0.3 mM NaGTP and 0.2 mM QX-314). Before use, internal solutions were filtered with a 0.22 μm syringe filter (Merck Millipore). Pipettes were mounted to a headstage (CV-7b, Molecular Devices, USA) and controlled via a Multiclamp 700B amplifier (Axon CNS, Molecular Devices). Following entry into whole cell configuration, access resistance (R_a_) was monitored every 2 minutes and experiments were only included if R_a_ remained stable and below 25 MΩ. Recordings were low-pass filtered online at 2 kHz (8-pole Bessel), acquired using Clampex software (pClamp 10, Molecular Devices), and exported into Matlab (R2017a, Mathworks) for off-line analysis using custom-made scripts.

To examine the direct effects of activating excitatory DREADDs upon interneuron excitability, current clamp recordings were conducted in aCSF containing kynurenic acid (3 mM) and hM_3_D_q_ receptors were activated by bath application of CNO (10-20 μM, Tocris, Bio-Techne). The spontaneous action potential firing rate of each interneuron was compared for a 5-minute period before and after CNO application, having allowed 3 minutes for the CNO to reach the chamber. To measure the post-synaptic GABAergic currents induced by activating hM_3_D_q_ receptors in a specific interneuron population, voltage-clamp recordings were conducted by clamping CA1 and CA3 pyramidal neurons at the reversal potential for glutamatergic current (E_GLUT_) in the presence of kynurenic acid. Once recordings had stabilised, the amplitude of post-synaptic inhibitory conductances were compared across two-minute periods recorded under baseline conditions, after bath application of CNO and then after co-administration of CNO and tetrodotoxin (TTX, 1-2 μM).

### Quantification of seizure-like events in vitro

A semi-automated detection algorithm was used to identify the start and end of individual spontaneous seizure-like events (SLEs) in vitro. Current-clamp traces were down-sampled to 1 kHz and then band-pass filtered (typically 0.05-to-0.2 Hz) using a Bessel filter (2^nd^ order). The signal was corrected for the rise time of the filter and subsequently rectified, thresholded and binarised, merging events that were close in time (typically under 1 minute apart), and ignoring events shorter than 5 seconds. Experiments to test the effects of a drug (e.g. CNO) comprised a 15-minute baseline, followed by a 3 minute period to allow the drug to reach the recording chamber, and a further 15-minute period in which the slice was continuously superfused with drug-containing aCSF. The 15-minute time periods (‘baseline’ and ‘drug’) were assessed using the same SLE detection settings. Total SLE activity was defined as the sum of time that the slice displayed SLE activity during a 15-minute period. SLE frequency was calculated from the number of SLEs that initiated during a 15-minute period and SLE length was the mean duration of individual SLEs that were completely contained within a 15-minute period.

### Viral transduction of hippocampal interneurons in vivo

All animal procedures relating to in vivo experiments were carried out with the approval of the local ethics committee for animal research in Bucharest and in accordance with European Union Directive 2010/63/EU on the protection of animals used for scientific purposes. For viral injections, adult animals of either sex were anaesthetised with isoflurane (maintained at 1.5-2%, 0.4 L/min) and mounted on a stereotaxic instrument (Kopf, RWD Life Science). The level of anaesthesia was continuously monitored, eye drops (Corneregel, Bausch & Lomb) were applied to avoid corneal desiccation and a heat pad system (DC Temperature Controller, FHC) was used to maintain the body temperature in the physiological range. Wiretrol II glass capillaries (Drummond Scientific) were pulled using a vertical puller (Narishige PC-10, Japan) and connected to the Hamilton syringe via compression fittings (RN 1 mm, Hamilton, USA). Small craniotomies were generated with a precision drill (FBS 240/E, Proxxon Micromot) and hippocampal bilateral injections were performed both dorsally (−2.18 anteroposterior (AP), 2.3 mediolateral (ML), 2.4-1.85 dorsoventral (DV), relative to Bregma) and ventrally (−2.7 AP, 2.9 ML, 3.1-2.25 DV, relative to Bregma) at a rate of 1.66 nL/s, slowly retreating the injection pipette (0.91 µm/s) to maximise delivery throughout the hippocampus. The injection was controlled via a micromanipulator (NeuroCraft MCM, FHC) attached to a syringe (705RN, 50 µL, Hamilton, USA). Each of four injection tracks was infused with 1350 nL of virus (Addgene, USA), of which 170 nL were delivered at each end of the track. After infusing the target volume of viral solution, a time window of 5 minutes allowed the virus solution to spread through the hippocampal tissue before the injection pipette was completely retracted. After allowing 2 to 6 months for expression, and at least three days before commencing seizure experiments, mice were anaesthetised and implanted with an infusion cannula (C315GS-5 guide cannula, Plastics One, USA) directly over the viral injection site in the right dorsal hippocampus. The cannula was secured to the skull via bone cement (Refobacin R40, Biomet UK).

### Quantification of epileptic behaviour in vivo

For each seizure experiment, mice were briefly anaesthetised with isoflurane to allow insertion of the infusion cannula into the guide cannula, such that the tip of the infusion cannula was located within the hippocampus at −2.18 AP, 2.3 ML, 2.2 DV, relative to Bregma. At this point, an intraperitoneal (i.p.) bolus of solution containing CNO (4 mg/kg with 4% dimethyl sulfoxide (DMSO) in saline) or vehicle (4% DMSO in saline) was administered and the mouse was allowed to fully recover from anaesthesia for 15 minutes before starting experiments. To monitor behaviour, the mouse was placed in a square arena (400 mm x 400 mm) in which it was able to move freely virtue of a connector assembly (C313C, Plastics One, USA) and swivel system (375/22PS blue, 22ga, Instech, USA), which connected the infusion tubing to a 1 µL syringe (7101, Hamilton, USA) controlled by an infusion pump (IVAC P6000, Cardinal Health). Following a 20 minute baseline period, 4-aminopyridine (4-AP) was infused directly into the hippocampus according to a spaced delivery protocol (three 4-AP infusions, each separated by 12 minutes and consisting of a 200 nL injection of 2 mM 4-AP over 2 minutes). Infusions were terminated immediately if the mouse reached the stage of generalised motor convulsions. Throughout each experiment video recordings were performed using two high speed, high definition cameras located at a right angle from each other (Hero 3+ Silver, GoPro, USA) at 60 frames per second, 1920×1080 pixels per frame. Polarised filters were used to reduce glare from the arena walls. A third camera captured the animals’ movements directly from above, allowing to track the location and locomotor activity of the mouse at any given time point. The epileptic behaviour was blindly scored using the Racine scale. Individual behaviours were considered as binary point events across time at a sampling frequency of 1 Hz. Events were then weighted according to the Racine classification of that behaviour: 1. orofacial clonic activity (Racine 1); 2. head nodding (Racine 2); 3. limb clonic activity (Racine 3); 4. retreating/rearing with orofacial clonic activity (Racine 4); 5. rearing and falling and/or jumping (Racine 5-generalised motor convulsions). Rearing and falling was considered as the ‘threshold seizure’ behaviour.

### Immunohistochemistry and quantification of interneuron distribution

For in vitro studies, organotypic hippocampal slices expressing hM_3_D_q_-mCherry or EGFP in specific interneuron populations were fixed overnight at 4°C in 4% paraformaldehyde with 4% sucrose, in 0.01 M phosphate buffer solution (PBS), pH 7.4. The slices used for immunofluorescence were washed and embedded in 3% agar, and re-sectioned at 50 μm on a vibrating microtome (Microm HM 650V, Thermo Fisher Scientific). For in vivo studies, mice expressing hM_3_D_q_-mCherry were transcardially perfused and the brains sectioned at 50 μm. PV expression was visualised by incubating sections in 1:500 guinea pig primary antibody (cat. no. 195 004, Synaptic Systems) in PBS with 0.3% Triton-X (PBST) with 1% normal goat serum (NGS, Thermo Fisher Scientific) overnight at 4°C, followed by 1:500 Alexa 488 goat anti-guinea pig secondary antibody (Thermo Fisher Scientific) in PBST with 1% NGS overnight at 4°C. For SST immunolabeling, the tissue was processed with a basic antigen retrieval kit at 95°C for 10 minutes (R&D Systems). All sections were pre-incubated in 10% NGS in PBST for at least 2 h at room temperature. SST expression was visualised by incubating sections in 1:250 rat primary antibody (MAB 354, Millipore) in PBST with 1% NGS for 11 days at 4°C, followed by 1:500 Alexa 488 goat anti-rat secondary antibody (Thermo Fisher Scientific) in PBST with 1% NGS for 2 days at 4°C. VIP expression was visualised by incubating sections in 1:5000 rabbit primary antibody (donation from Professor Peter P. Somogyi) in PBST with 1% NGS for 2 days at 4°C, followed by 1:500 Alexa 488 goat anti-rabbit secondary antibody (Thermo Fisher Scientific) in PBST with 1% NGS overnight at 4°C. Finally, all sections were mounted in Vectashield (Vector Laboratories) and images were captured with an LSM 880 confocal microscope equipped with 488 nm and 561 nm lasers, a 20x water-immersion objective (W Plan-Apochromat, NA 1.0) and controlled via the ZEN black software (Zeiss).

To determine the number and distribution of soma and processes associated with each interneuron subtype, we performed quantitative image analysis on organotypic hippocampal slices from PV-cre, SST-cre and VIP-cre mice that had been injected with floxed AAVs. Slices were subjected to confocal microscopy and, in the resulting images, the CA areas were linearized along the pyramidal layer and the soma of fluorescent neurons were automatically detected via a custom-made, two-pass algorithm extracting maximally stable extremal region features using Matlab (Matas et al., 2004; Nistér and Stewénius, 2008). The number of virally-transduced interneurons was derived directly from the number of fluorescent soma per optical section. Meanwhile, to describe the distribution of processes associated with each interneuron subtype, soma were digitally removed from the linearized images to generate a transverse expression profile of fluorescent processes relative to the pyramidal cell layer. To compare across multiple interneuron populations, these expression profiles were normalised by the area under each curve.

### Data analysis

Digital signal processing and presentation were performed using custom-made programs in the Matlab environment (R2017b, Mathworks). Figures were built using vector-based graphic design in CorelDraw (X6, Corel Corporation) and the statistical software GraphPad Prism (v6.01, GraphPad Software). Video data processing for tracking locomotor activity of animals was performed using the open-source software Bonsai v2.3 (Lopes et al., 2015). Data are presented as mean ± standard error of mean (SEM) and the statistical tests are reported at the relevant points in the text (GraphPad Prism; Matlab). Non-parametric tests were used when a normal distribution of data could not be ascertained. Appropriate post-hoc tests were used when ANOVA tests confirmed a statistically significant effect.

## Results

### The activity of distinct hippocampal GABA-releasing interneuron populations can be enhanced with excitatory DREADDs

To examine the potential of chemicogenetically enhancing the synaptic output of hippocampal interneurons, we used mouse organotypic hippocampal brain slices. This system enabled us to perform targeted patch clamp recordings to determine the pre-synaptic and post-synaptic efficacy of DREADDs, as well as the opportunity to examine the impact of interneuron recruitment upon spontaneously generated epileptiform activity. As in previous work (Trevelyan et al., 2006; Fujiwara-Tsukamoto et al., 2007; Sessolo et al., 2015), our recordings from excitatory pyramidal neurons within the Cornu Ammonis (CA) areas revealed that organotypic hippocampal brain slices exhibit reproducible spontaneous SLEs that can be quantified and exhibit a sustained duration (mean duration was 48.3±8 s) and stable frequency (**Fig. 1***A,B*; Dyhrfjeld-Johnsen et al., 2010; Berdichevsky et al., 2012; Lillis et al., 2015; Liu et al., 2017). Consistent with epileptiform activity in many systems (Trevelyan et al., 2006; Fujiwara-Tsukamoto et al., 2007; Sessolo et al., 2015), these spontaneous SLEs recruited both excitatory and inhibitory neurons within the hippocampal network. Paired recordings revealed that during each SLE, both excitatory and inhibitory neurons are recruited within the CA areas (**Fig. 1***C*), which results in intense barrages of glutamatergic and GABAergic post-synaptic currents converging upon pyramidal neurons (**Fig. 1***D*). By combining promoter-specific cre-recombinase mice with floxed chemicogenetic constructs, we were then able to investigate the cellular and network effects of delivering DREADDs to specific interneuron populations.

**Fig. 1.**
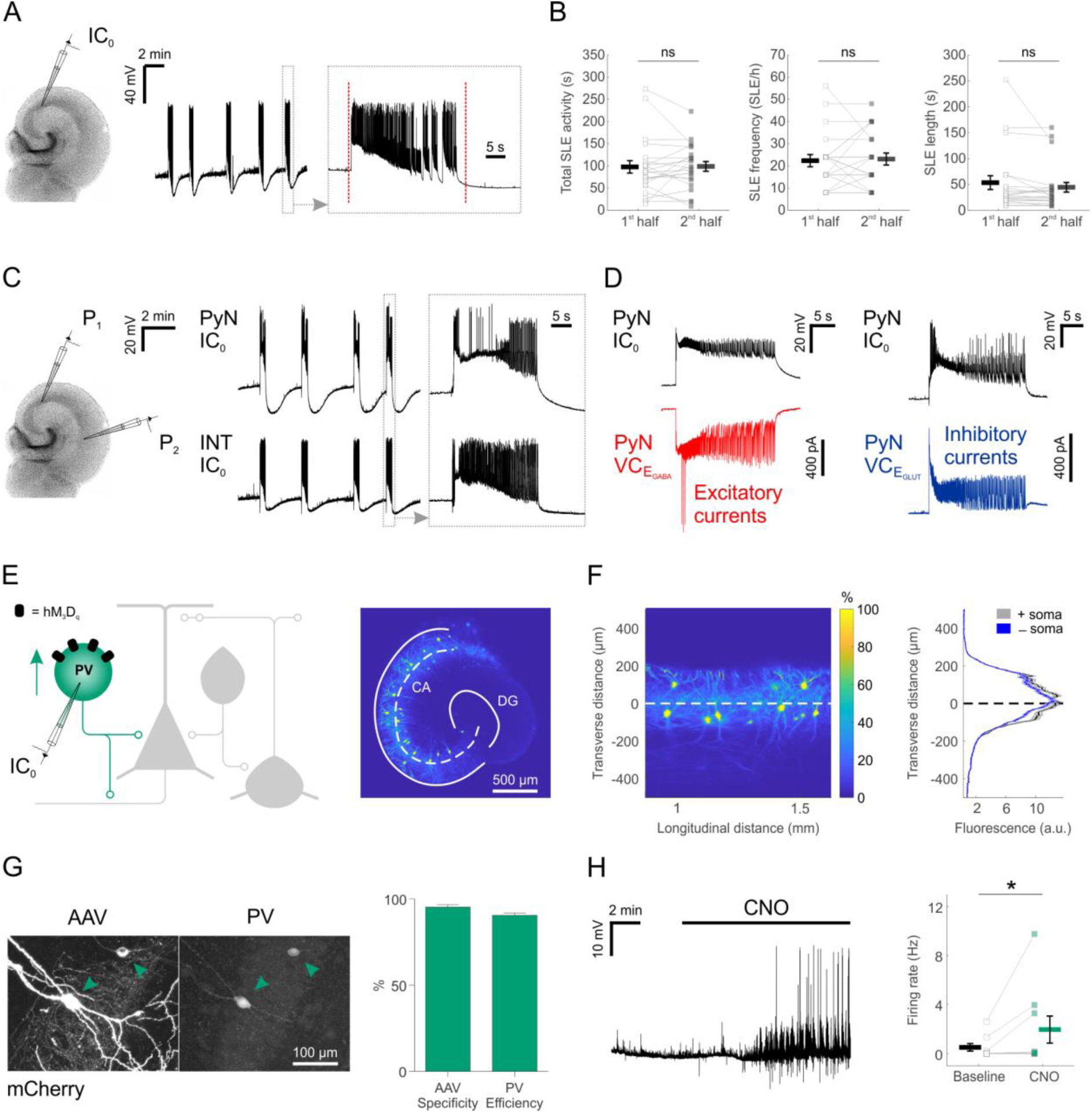
Chemicogenetic recruitment of hippocampal interneurons in an in vitro model of epileptiform activity. ***A***, Current-clamp recording of a CA3 pyramidal neuron from a mouse organotypic hippocampal slice reveals repeated SLEs. The vertical red dotted lines in the expanded trace mark the onset and the termination of the underlying SLE, as determined by an automated detection algorithm. ***B***, A 15-minute time window was used to assess the stability of the epileptiform activity. Total SLE activity, defined as the cumulative duration of SLE activity (left, 96.8±13.9 s during 1^st^ half, 97.9±10.9 s during 2^nd^ half; N=22 slices, W_(21)_=110.5, p=0.8715, two-tailed Wilcoxon signed-rank test), SLE frequency (middle, 22.2±2.7 SLEs/h during 1^st^ half, 22.9±2.7 SLEs/h during 2^nd^ half; N=22 slices, W(12)=34, p=0.7153, two-tailed Wilcoxon signed-rank test) and individual SLE length (right, 52.8±13.4 s during 1^st^ half, 43.9±9.2 s during 2^nd^ half; N=22 slices, W_(21)_=71, p=0.1281, two-tailed Wilcoxon signed-rank test) were stable across the 15-minute time window. ***C***, A dual current-clamp (IC0) electrophysiological recording of a CA_3_ pyramidal neuron (PyN) and a GABAergic interneuron (INT) reveal spontaneous SLEs in a mouse organotypic hippocampal brain slice (left). Expanded traces (right) show that both excitatory and inhibitory neurons are recruited during the SLEs. ***D***, Simultaneous current clamp and voltage clamp (VC) recordings from pairs of pyramidal neurons demonstrate that strong barrages of excitatory (bottom-left, red) and inhibitory (bottom-right, blue) post-synaptic currents occur throughout the SLEs, as monitored by current clamp recordings from a neighbouring pyramidal neuron (top). EGABA, reversal potential for GABAergic current; EGLUT, reversal potential for glutamatergic current. ***E***, Cartoon of hippocampal circuitry (left) showing the targeting of the hM_3_D_q_ receptor to PV interneurons. Confocal image (right) of a hippocampal slice from a PV-cre mouse illustrates the fluorescence distribution profile of virally-transduced PV interneurons. The superimposed white dashed line marks the centre of the pyramidal cell layer. Continuous white lines indicate the dentate gyrus (DG) and outline of the CA areas (CA). ***F***, The confocal image was linearized (left) to facilitate the quantification of the transverse expression profile for PV interneurons, relative to the pyramidal cell layer (dashed line at zero). This confirmed that PV interneurons (+soma) and their processes (-soma) were restricted to the pyramidal cell layer (right). ***G***, The immunohistochemical characterisation of PV interneurons transduced with AAV_8_-hSyn-DIO-hM_3_D_q_-mCherry demonstrates high targeting specificity and efficiency (N=14 sections from 6 slices). ***H***, An example current-clamp recording of a PV interneuron expressing hM_3_D_q_ receptors shows that CNO promoted action potential firing (left). Recordings were conducted in kynurenic acid to isolate the direct effects of hM_3_D_q_ receptor activation. Population data shows a significant increase in firing rate following CNO (right; N=9 slices, W_(7)_=0, p=0.0156, two-tailed Wilcoxon signed-rank test)

PV interneurons are a principal GABAergic population of the hippocampus, which reside primarily within the pyramidal layer of all CA areas (Pawelzik et al., 2002; Klausberger and Somogyi, 2008), and whose axons target the somatic compartment of pyramidal neurons (Pawelzik et al., 2002; Bartos and Elgueta, 2012; Hu et al., 2014). Consistent with this, injection of AAV containing floxed constructs into organotypic hippocampal slices from PV-cre mice resulted in somatic and process expression that was restricted to the pyramidal cell layer (**Fig. 1***E,F*). This is consistent with previous observations that organotypic slices retain fundamental features of the intact circuit, including the subcellular targeting of perisomatic and dendritic domains of pyramidal neurons by distinct interneuron populations (Streit et al., 1989; De Simoni et al., 2003; Cristo et al., 2004). Our immunohistochemical experiments confirmed that the PV interneurons could be efficiently and specifically targeted with a floxed excitatory DREADD construct, which resulted in the expression of the human type-3 muscarinic designer receptors coupled with the G_q_ protein (hM_3_D_q_ receptor). Two-to-four weeks after viral transduction with AAV_8_-hSyn-DIO-hM_3_D_q_-mCherry, the majority of expressing neurons were shown to be immunopositive for PV (95.2±1.5% ‘specificity’) and the majority of all PV immunopositive neurons were expressing hM_3_D_q_-mCherry (90.5±1.4% ‘efficiency’; **Fig. 1***G*). To assess whether the hM_3_D_q_ receptors could be activated and increase the output of the PV interneuron population, current clamp recordings were targeted to hM_3_D_q_-mCherry positive neurons. The addition of the hM_3_D_q_ ligand, CNO, led to a significant increase in the firing rate of PV interneurons, from 0.5±0.3 Hz during baseline, to 1.9±1.1 Hz in the presence of CNO (**Fig. 1***H*).

To examine the potential to chemicogenetically enhance the output of two other major hippocampal interneuron subtypes, we used organotypic hippocampal slices generated from SST-cre mice and VIP-cre mice, in order to target hM_3_D_q_ receptors to SST and VIP interneurons, respectively (**Fig. 2**). Consistent with previous evidence that SST interneurons target the dendritic compartments of hippocampal principal neurons (Katona et al., 1999; Lovett-Barron et al., 2012), we confirmed that the soma and processes of SST interneurons are located within stratum oriens and lacunosum-moleculare, and tend to avoid the pyramidal cell layers of the CA regions (**Fig. 2***A,B*). Immunohistochemistry confirmed that the SST interneurons could be efficiently and specifically targeted with hM_3_D_q_ receptors in SST-cre slices. Two-to-four weeks after viral transduction with AAV_8_-hSyn-DIO-hM_3_D_q_-mCherry, the majority of expressing neurons were immunopositive for SST (94.1±1.6 specificity) and the majority of all SST immunopositive neurons were expressing hM_3_D_q_-mCherry (89.6±1.5% efficiency; **Fig. 2***C*). Finally, to assess whether the hM_3_D_q_ receptors could be activated and increase the output of the SST interneuron population, current clamp recordings were targeted to hM_3_D_q_-mCherry positive neurons. The addition of CNO was shown to cause a significant increase in the firing rate of SST interneurons, from 1.1±0.5 Hz during baseline, to 3.7±1.3 Hz in the presence of CNO (**Fig. 2***D*).

**Fig. 2.**
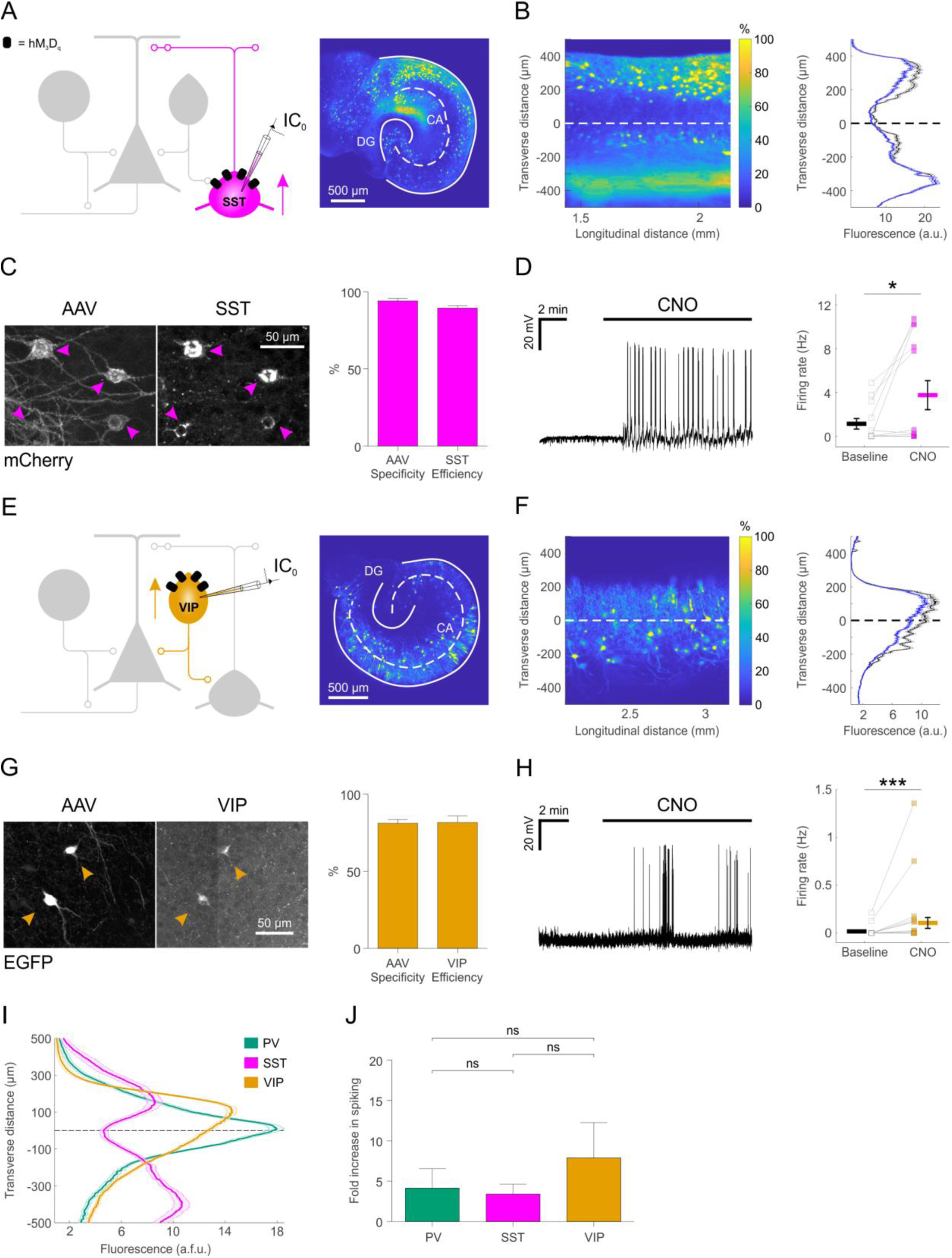
Distinct subtypes of hippocampal GABAergic interneurons can be recruited ^via excitatory DREADDs. ***A***, Cartoon (left) showing the targeting of hM_3_D_q_ receptors to SST hippocampal interneurons. A confocal image of a SST-cre mouse slice (right) illustrates the fluorescence distribution profile of virally-transduced SST interneurons. Continuous white lines outline the DG and CA areas. ***B***, The confocal image was linearized (left) to facilitate quantification of the transverse expression profile for SST interneurons, relative to the pyramidal cell layer (dashed white line at zero). This confirmed that SST interneurons (+soma) and their processes (-soma) were associated with stratum oriens and lacunosum-moleculare (right). ***C***, Immunohistochemical characterisation of SST interneurons transduced with AAV8-hSyn-DIO-hM_3_D_q_-mCherry demonstrates high targeting specificity and efficiency (N=16 sections from 6 slices). ***D***, Current-clamp recording from a SST interneuron ^expressing hM_3_D_q_ in the presence of kynurenic acid, showing that CNO promotes^ action potential firing (left). Population data shows a significant increase in firing ^rate in the presence of CNO (right, N=13 slices, W(9)=4, p=0.0273, two-tailed Wilcoxon signed-rank test). ***E***, Cartoon (left) shows the targeting of hM_3_D_q_ receptors^ to VIP interneurons and confocal image (right) illustrates the fluorescence distribution profile of virally-transduced interneurons in a VIP-cre slice. ***F***, Linearizing the confocal image (left) confirmed that the expression profile for VIP interneurons (+soma) and their processes (-soma) was associated with stratum radiatum/lacunosum-moleculare, pyramidale and oriens (right). ***G***, ^Immunohistochemical characterisation of VIP interneurons transduced with AAV8-hSyn-DIO-hM_3_D_q_-mCherry demonstrates high targeting specificity and efficiency^ (N=4 sections from 2 slices). ***H***, Example current-clamp recording from a VIP interneuron expressing hM_3_D_q_ in kynurenic acid, showing that CNO promotes action potential firing (left). Population data (right) shows a significant increase in firing rate in the presence of CNO (N=27 slices, W(13)=1, p=0.0005, two-tailed Wilcoxon^ signed-rank test). ***I***, Comparison of the fluorescence distribution profiles following viral-transduction of the three interneuron populations (PV, N=38 slices; SST, N=37 slices; VIP, N=57 slices). Each distribution was normalised by the area under the profile curve and shown to be significantly different between the cell types (interaction between cell type and location: F_(8,516)_=58.81; p<0.0001, repeated measures two-way ANOVA). ***J***, There was no difference in the CNO-induced fold-increase in spiking rate (normalised to baseline) between the three interneuron populations (χ2_(2)_ =0.9136, p=0.6333, Kruskal-Wallis test).

We next assessed the potential to chemicogenetically target the VIP interneuron population (**Fig. 2***E*). Consistent with previous evidence showing that distinct VIP interneuron subtypes target O-LM interneurons and the perisomatic regions or proximal dendrites of pyramidal neurons (Léránth et al., 1984; Acsády et al., 1996; Chamberland and Topolnik, 2012), we confirmed that the soma and processes of virally-transduced VIP interneurons were located within stratum radiatum/lacunosum-moleculare, pyramidale and oriens in slices from the VIP-cre mice (Köhler, 1982; Taniguchi et al., 2011; Botcher et al., 2014) (**Fig. 2***F*). Immunohistochemistry confirmed that the VIP interneurons could be efficiently and specifically targeted with floxed constructs delivered by AAV. Two-to-four weeks after transduction, the majority of expressing neurons were immunopositive for VIP (81.2±2.4% specificity) and the majority of VIP immunopositive neurons expressed the floxed construct (81.6±4.4% efficiency; **Fig. 2***G*). Finally, activating hM_3_D_q_ receptors targeted to VIP interneurons by CNO increased the firing rate from 0.01±0.01 Hz during baseline, to 0.1±0.1 Hz in the presence of CNO (**Fig. 2***H*).

A summary plot for the three interneuron populations confirmed that the expression profiles of PV, SST and VIP interneurons were significantly different from one another (**Fig. 2***I*) and were consistent with data from acute preparations of mouse hippocampus (Taniguchi et al., 2011; Lovett-Barron et al., 2012). Meanwhile, CNO-mediated activation of hM_3_D_q_ receptors resulted in comparable fold-increases in spiking activity across the three interneuron populations: PV interneurons showed a 4.1±2.4 fold increase in their firing rate, SST interneurons showed a 3.4±1.2 fold increase and VIP interneurons showed a 7.9±4.4 fold increase (**Fig. 2***J*).

### Chemicogenetic enhancement of specific GABAergic interneuron populations can attenuate hippocampal epileptiform activity in vitro

To investigate the potential anti-epileptic effects of chemicogenetically enhancing the output of interneuron populations, excitatory DREADDs were targeted to one of the interneuron subtypes in organotypic hippocampal slices from PV-cre, SST-cre or VIP-cre mice. After 2-to-4 weeks to allow for viral-mediated expression of the hM_3_D_q_ receptors, spontaneous SLEs were recorded from CA1 or CA3 pyramidal neurons before and after CNO-mediated activation of the relevant interneuron population. Automated detection of the spontaneous SLEs (see Materials and Methods) provided quantification of the total SLE activity and then a breakdown in terms of the effects upon SLE frequency and individual SLE duration. In PV-targeted slices (**Fig. 3***A*), CNO-mediated activation of hM_3_D_q_ receptors significantly reduced the total SLE activity (from 247.5±47.8 s during baseline to 116.5±47.4 s during CNO), which resulted from a significant decrease in SLE frequency (from 17.6±5.3 SLEs/h during baseline to 8.4±3.3 SLEs/h during CNO), without affecting the duration of individual SLEs (65.8±15.7 s during baseline and 77.5±27.4 s during CNO) (**Fig. 3***B*). In SST-targeted slices (**Fig. 3***C*), CNO-mediated activation of hM_3_D_q_ receptors also significantly reduced total SLE activity (from 263.2±41.4 s during baseline to 133.3±40.2 s during CNO), which involved a decrease in SLE frequency (from 26±6.8 SLEs/h during baseline to 13.3±3.5 SLEs/h during CNO), without significantly affecting individual SLE length (72±21.7 s during baseline and 47.6±14.5 s during CNO) (**Fig. 3***D*). In VIP-targeted slices (**Fig. 3***E*), CNO-mediated activation of hM_3_D_q_ receptors did not decrease the total SLE activity (234.3±40.6 s during baseline and 256.3±62.7 s during CNO). Whilst enhancing VIP interneuron output led to a decrease in SLE frequency (from 21.7±4.3 SLEs/h during baseline to 17.3±4 SLEs/h during CNO), there was a simultaneous significant increase in the duration of individual SLEs (from 43.5±10.3 s during baseline to 69.4±19.6 s during CNO) (**Fig. 3***F*). Finally, to assess potential off-target effects of CNO, the drug was bath-applied during the recording of spontaneous SLEs in control slices that were not expressing excitatory DREADDs (**Fig. 3***G*). In these experiments, no changes were detected for total SLE activity (329.9±63.8 s during baseline and 332.8±82.3 s during CNO), SLE frequency (23.2±5.8 SLEs/h during baseline and 20±5.1 SLEs/h during CNO), or individual SLE length (56.5±12.4 s during baseline and 46.2±12.2 s during CNO) (**Fig. 3***H*).

**Fig. 3.**
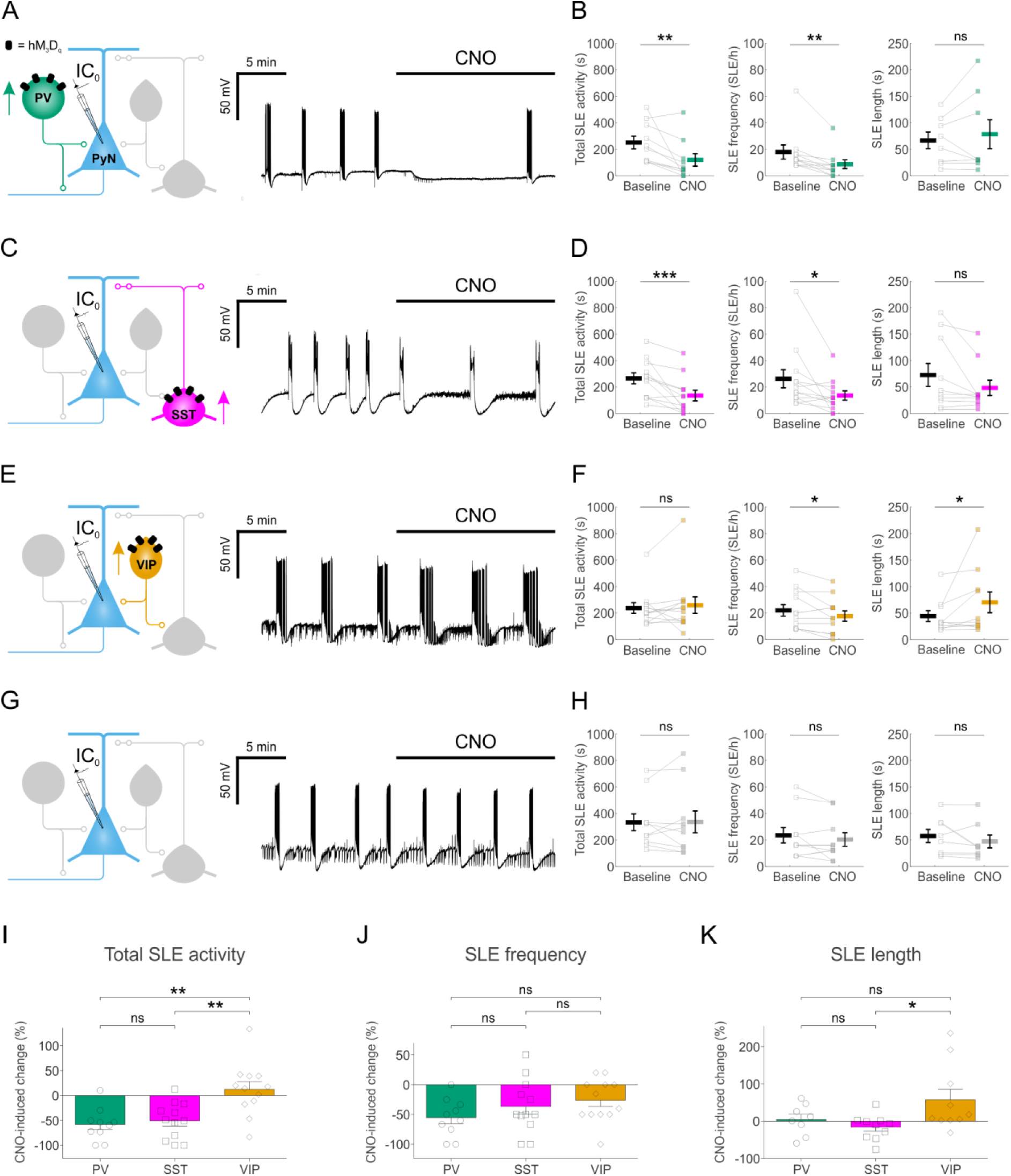
Chemicogenetic enhancement of specific GABAergic interneuron populations attenuates hippocampal SLEs. ***A***, PV-cre mice and floxed viral constructs were used to target hM_3_D_q_ receptors to PV interneurons in organotypic hippocampal brain slices. The effects of CNO upon spontaneous SLEs were monitored by current-clamp recordings from CA1 or CA3 pyramidal neurons. ***B***, Population data from experiments targeting PV interneurons (N=10 slices) showed a reduction in total SLE activity (left, W_(10)_=1, p=0.0039, two-tailed Wilcoxon signed-rank test), a decrease in SLE frequency (middle, W_(9)_=0, p=0.0039, two-tailed Wilcoxon signed-rank test), and no change in SLE length (right, W_(8)_=12, p=0.4609, two-tailed Wilcoxon signed-rank test) following addition of CNO. ***C***, SST-cre mice were used to target hM_3_D_q_ receptors to SST interneurons. ***D***, Population data from experiments targeting SST interneurons (N=12 slices) showed a reduction in total SLE activity (left, W_(12)_=1, p=0.001, two-tailed Wilcoxon signed-rank test), a decrease in SLE frequency (middle, W_(12)_=6, p=0.0166, two-tailed Wilcoxon signed-rank test), and no change in SLE length (right, W(10)=10, p=0.0840, two-tailed Wilcoxon signed-rank test) following addition of CNO. ***E***, VIP-cre mice were used to target hM_3_D_q_ receptors to VIP interneurons. ***F***, Population data from experiments targeting VIP interneurons (N=12 slices) showed no change in total SLE activity (left, W_(12)_=25, p=0.3013, two-tailed Wilcoxon signed-rank test), a reduction in SLE frequency (middle, t_(11)_=2.24, p=0.0468, two-tailed paired t-Test), and an increase in SLE length (right, W_(10)_=6, p=0.0273, two-tailed Wilcoxon signed-rank test) following addition of CNO. ***G***, Control experiments were conducted on slices that had not received floxed DREADD constructs. ***H***, Population data (N=10 slices) demonstrated no change in total SLE activity (left, t_(9)_=-0.08, p=0.9337, two-tailed paired t-Test), SLE frequency (middle, t_(9)_=1.31, p=0.2229, two-tailed paired t-Test), or SLE length (right, W_(8)_=6, p=0.1094, two-tailed Wilcoxon signed-rank test) following the addition of CNO to control slices. ***I***, Total SLE activity was significantly reduced by chemicogenetically enhancing either PV or SST interneurons, compared to enhancing VIP interneurons (F_(2,31)_=9.747, p=0.0005, one-way ANOVA, followed by Tukey’s post-hoc multiple comparison tests; VIP vs PV, p=0.0013; VIP vs SST, p=0.0025). ***J***, The frequency of SLEs was reduced when PV interneurons, SST interneurons or VIP interneurons were targeted. No significant difference in the reduction in SLE frequency was detected across the three interneurons (F_(2,31)_=1.583, p=0.2215, one-way ANOVA). ***K***, Individual SLEs became significantly longer when VIP interneurons were recruited, compared to SST interneurons (F_(2,25)_=3.772, p=0.037, one-way ANOVA, followed by Tukey’s post-hoc multiple comparison tests; VIP vs SST, p=0.0337). * indicates p<0.05, ** indicates p<0.01.

Comparisons across the different interneuron populations confirmed subtype-specific effects. Total SLE activity was reduced by more than half following activation of either PV interneurons (down 58.2±10.3%) or of SST interneurons (down 50.8±10.5%), and both of these reductions were significantly greater than the change in total SLE activity induced by recruiting VIP interneurons (up 12.7±15.4%) (**Fig. 3***I*). Each of the interneuron populations was able to decrease SLE frequency: PV interneurons by 55.4±10.1%, SST interneurons by 36.7±13% and VIP interneurons by 26.3±10.4% (**Fig. 3***J*). Meanwhile, only the VIP population increased individual SLE length by 57.5±28.4%, as compared to the effect upon SLE length of activating SST (−16.1±10.6%) or PV interneurons (4.4±14.7%) (**Fig. 3***K*). These observations are consistent with previous evidence that VIP interneurons can actually promote ongoing epileptiform activity, perhaps because they mediate disinhibitory effects as a result of preferentially innervating other GABAergic interneurons (Pi et al., 2013; Karnani et al., 2016; Khoshkhoo et al., 2017; Ye and Kaszuba, 2017). In summary therefore, chemicogenetically enhancing the output of interneuron populations can generate effective anti-epileptic effects, but the effectiveness of this approach depends upon the interneuron subtype.

### Chemicogenetically-enhanced interneuron subtypes differ in their post-synaptic inhibition of pyramidal neurons

To examine the cellular basis of these anti-epileptic effects, we performed voltage clamp recordings to compare how chemicogenetic recruitment affects the post-synaptic inhibition that converges upon pyramidal neurons. CA1 and CA_3_ pyramidal neurons were recorded with a caesium-based internal solution containing QX-_314_, and held at E_GLUT_ to isolate inhibitory post-synaptic currents (**Fig. 4***A,C,E*). For each of the three interneuron populations (PV, SST and VIP), CNO-activation of hM_3_D_q_ receptors resulted in a significant increase in post-synaptic inhibitory input (**Fig. 4***A,C,E*). Furthermore, for each interneuron population, the CNO-induced increases in post-synaptic inhibition were abolished by bath application of the voltage-gated Na^+^ blocker, TTX (1-2 μM; **Fig. 4***A,C,E*). This confirmed that in each case, the post-synaptic inhibition was the result of CNO-mediated increases in action potential-evoked GABA release from the interneurons. Following recruitment of PV interneurons, the total inhibitory post-synaptic input to pyramidal neurons increased from 11±4.2 pA/ms to 34±6.1 pA/ms and was abolished by TTX (−0.5±0.1 pA/ms; **Fig. 4***B*). Following recruitment of SST interneurons, the total inhibitory post-synaptic input to pyramidal neurons increased from 9.3±2.6 pA/ms to 126.6±14.6 pA/ms, and was abolished by TTX (0.3±0.5 pA/ms; **Fig. 4***D*). Following recruitment of VIP interneurons, the total inhibitory post-synaptic input to pyramidal neurons increased from 5.7±1.1 pA/ms to 16.2±2.4 pA/ms, and was abolished by TTX (0.4±0.7 pA/ms; **Fig. 4***F*). At the population level, the overall increase in post-synaptic inhibitory input resulting from CNO-mediated activation of hM_3_D_q_ receptors was highest for the SST interneuron population (increase of 116.7±15.5 pA/ms), then the PV interneuron population (increase of 22.6±5.7 pA/ms), and then the VIP interneuron population (increase of 10.3±2.3 pA/ms).

**Fig. 4.**
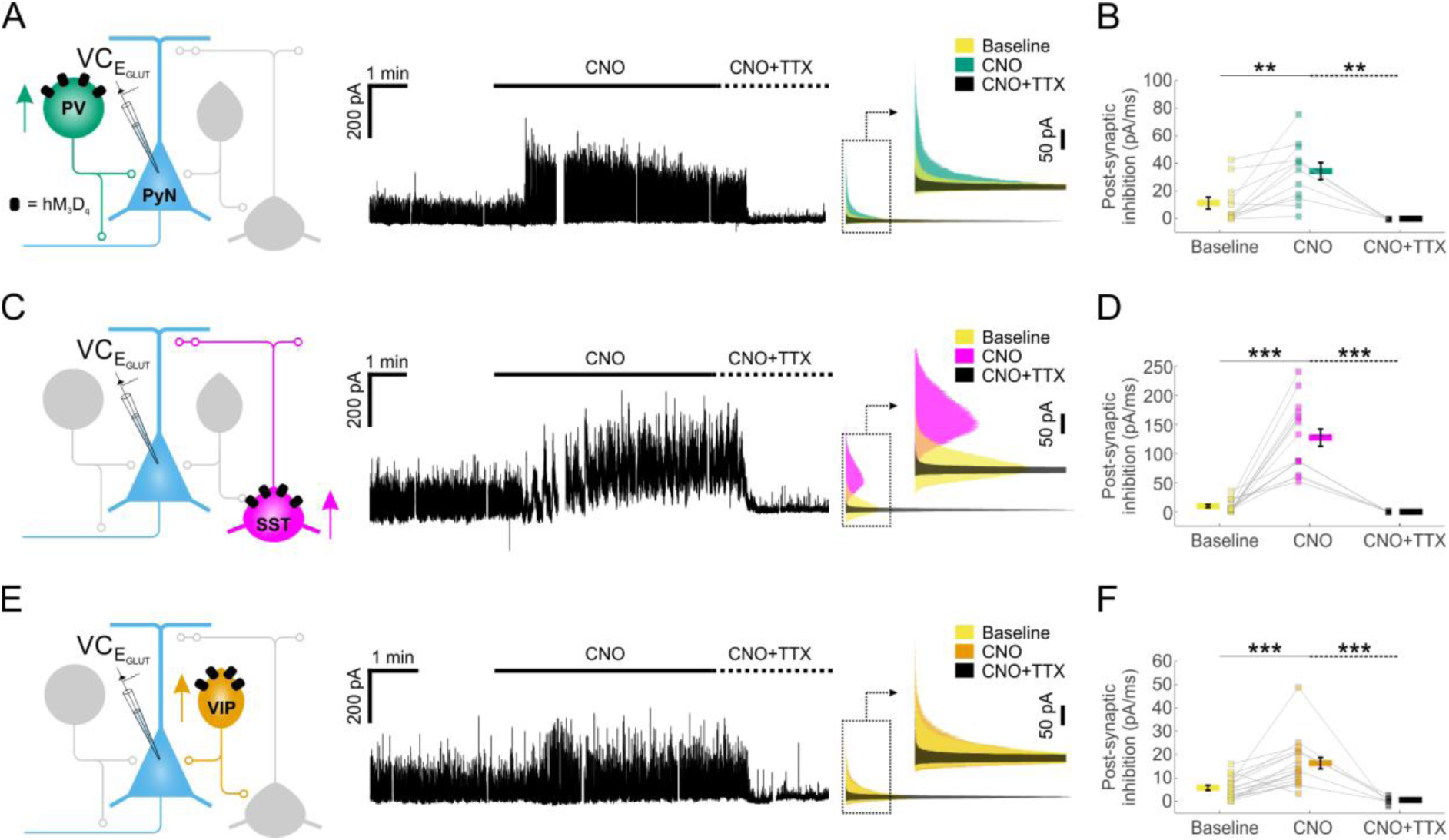
Chemicogenetic recruitment of interneuron populations generates different ^amounts of post-synaptic inhibition in pyramidal neurons. ***A***, hM_3_D_q_ receptors were targeted to PV interneurons and voltage-clamp recordings (at EGLUT) were performed^ from pyramidal neurons (left). CNO application elicited a pronounced increase in inhibitory post-synaptic currents converging upon the pyramidal neuron (middle). This increase was associated with spiking activity as it was abolished by bath application of TTX. Overlapping histograms (right) illustrate the probability distribution functions for the inhibitory post-synaptic currents during baseline, CNO, and co-administration of CNO plus TTX. ***B***, PV recruitment resulted in a significant change in total post-synaptic inhibitory charge converging onto pyramidal neurons ^(F(2,26)=9.541, p=0.0008, one-way ANOVA, followed by post-hoc Sidak’s multiple^ comparisons tests; baseline vs. CNO, N=12, p=0.0047; CNO vs. CNO plus TTX, N=5, ^p=0.0013). ***C***, hM_3_D_q_ receptors were targeted to SST interneurons and all^ conventions are the same as in ‘A’. ***D***, SST recruitment resulted in a significant change in total post-synaptic inhibitory charge converging onto pyramidal neurons (χ2_(2)_ =29.77, p<0.0001, Kruskal-Wallis test, followed by post-hoc Dunn’s multiple comparisons tests; baseline vs. CNO, N=16, p<0.0001; CNO vs. co-administration of CNO plus TTX, N=5, p<0.0001). ***E***, hM_3_D_q_ receptors were targeted to VIP interneurons and all conventions are the same as in ‘A’. ***F***, VIP recruitment resulted in a significant change in total post-synaptic inhibitory charge converging onto pyramidal neurons (F_(2,39)_=14.09, p<0.0001 by one-way ANOVA, followed by post-hoc Sidak’s multiple comparisons tests; baseline vs. CNO, N=18, p=0.0003; CNO vs. co-administration of CNO plus TTX, N=6, p=0.0001). ** indicates p<0.01, *** indicates p<0.001.

To compare across the different subtypes, we expressed our measurements of inhibitory efficacy in terms of individual interneurons. First, the CNO-induced increase in post-synaptic inhibitory input was normalised by the number of hM_3_D_q_-expressing cells, as determined from stereological cell counts in the hippocampal slices (**Fig. 5***A*; see Materials and Methods). The amount of inhibitory post-synaptic input elicited by CNO activation was found to be similar for an individual PV interneuron (1.0 ± 0.3 fold, relative to a PV interneuron) and an SST interneuron (1.1 ± 0.1 fold, relative to a PV interneuron), both of which were five times greater than for an individual VIP interneuron (0.2 ± 0.1 fold, relative to a PV interneuron; **Fig. 5***B*). Second, to relate this to anti-epileptic efficacy, we normalised the effects upon total SLE activity by the number of hM_3_D_q_-expressing cells. These data indicated that individual PV interneurons had the greatest anti-epileptic effect (1.0 ± 0.2 fold, relative to a PV interneuron), which was at least five times more than the anti-epileptic effect associated with an individual SST interneuron (0.2±0.04 fold, relative to a PV interneuron) or VIP interneuron (−0.1±0.1 fold, relative to a PV interneuron; **Fig. 5***C*). These observations are consistent with the idea that, as a result of their peri-somatic targeting of pyramidal neurons and their extensive axonal arbors, the recruitment of an individual PV interneuron can mediate particularly effective inhibition of pyramidal neurons (Freund and Buzsáki, 1996; Miles et al., 1996; Pouille et al., 2013).

**Fig. 5.**
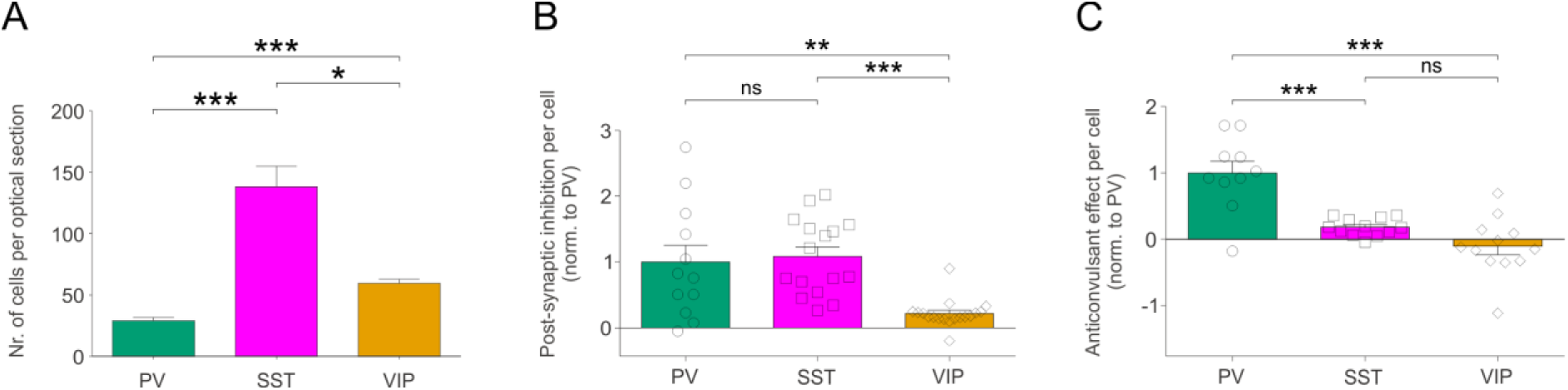
Chemicogenetically-enhanced interneuron subtypes differ in their post-synaptic inhibition of hippocampal pyramidal neurons. ***A***, The average number of hM_3_D_q_ expressing interneurons (per 24 μm optical section) differs significantly between the three interneuron populations, with an average of 29.1±2.7 PV cells, 138.1±16.8 SST interneurons, and 59.7±3.2 VIP cells (χ^2^(2)=54.44, p<0.0001, Kruskal-Wallis test, followed by Dunn’s multiple post-hoc comparisons; PV vs SST, p<0.0001; PV vs VIP, p<0.0001; SST vs VIP, p=0.0118). ***B***, Normalizing by the size of each interneuron population (i.e. number of hM_3_D_q_-expressing cells per slice), PV and SST interneurons were associated with similar amounts of post-synaptic inhibition, and both were significantly greater than that associated with VIP interneurons (χ_2_(2)=20.46, p<0.0001, Kruskal-Wallis test, followed by Dunn’s multiple post-hoc comparisons; PV vs SST, p>0.9999; PV vs VIP, p=0.0064; SST vs VIP, p<0.0001). ***C***, Normalizing by the size of each interneuron population, PV interneurons had the greatest effect upon the total SLE activity (shown in **Fig. 3***I*) (F_(2,31)_=21.03, p<0.0001, one-way ANOVA, followed by post-hoc Bonferroni’smultiple comparisons tests; PV vs SST, p=0.0002; PV vs VIP, p<0.0001; SST vs VIP, p=0.2808). * indicates p<0.05, ** indicates p<0.01, *** indicates p<0.001.

### Chemicogenetic recruitment of PV interneurons attenuates seizure activity in vivo

To further assess the potential of this chemicogenetic strategy, we designed an in vivo study to investigate the anti-epileptic effects of enhancing interneuron output. Given our findings in vitro, the PV interneurons were selected as an effective cell population to target with excitatory DREADDs in a temporal lobe model of in vivo seizure activity. The hM_3_D_q_ receptor was delivered to PV hippocampal interneurons by bilateral injections of AAV_8_-hSyn-DIO-hM_3_D_q_-mCherry in the hippocampus of 4-to-14-month-old PV-cre mice (**Fig. 6***A*). The virus was delivered at multiple depths in the ventral and dorsal hippocampus, which resulted in extensive expression of the hM_3_D_q_ receptor across the rostro-caudal axis (**Fig. 6***B,C*). The majority of all virally-transduced neurons were immunopositive for PV (86.42 ± 1.43% specificity) and the majority of PV immunopositive neurons expressed hM_3_D_q_-mCherry (89.17±2.12% efficiency; **Fig. 6***D*).

**Fig. 6.**
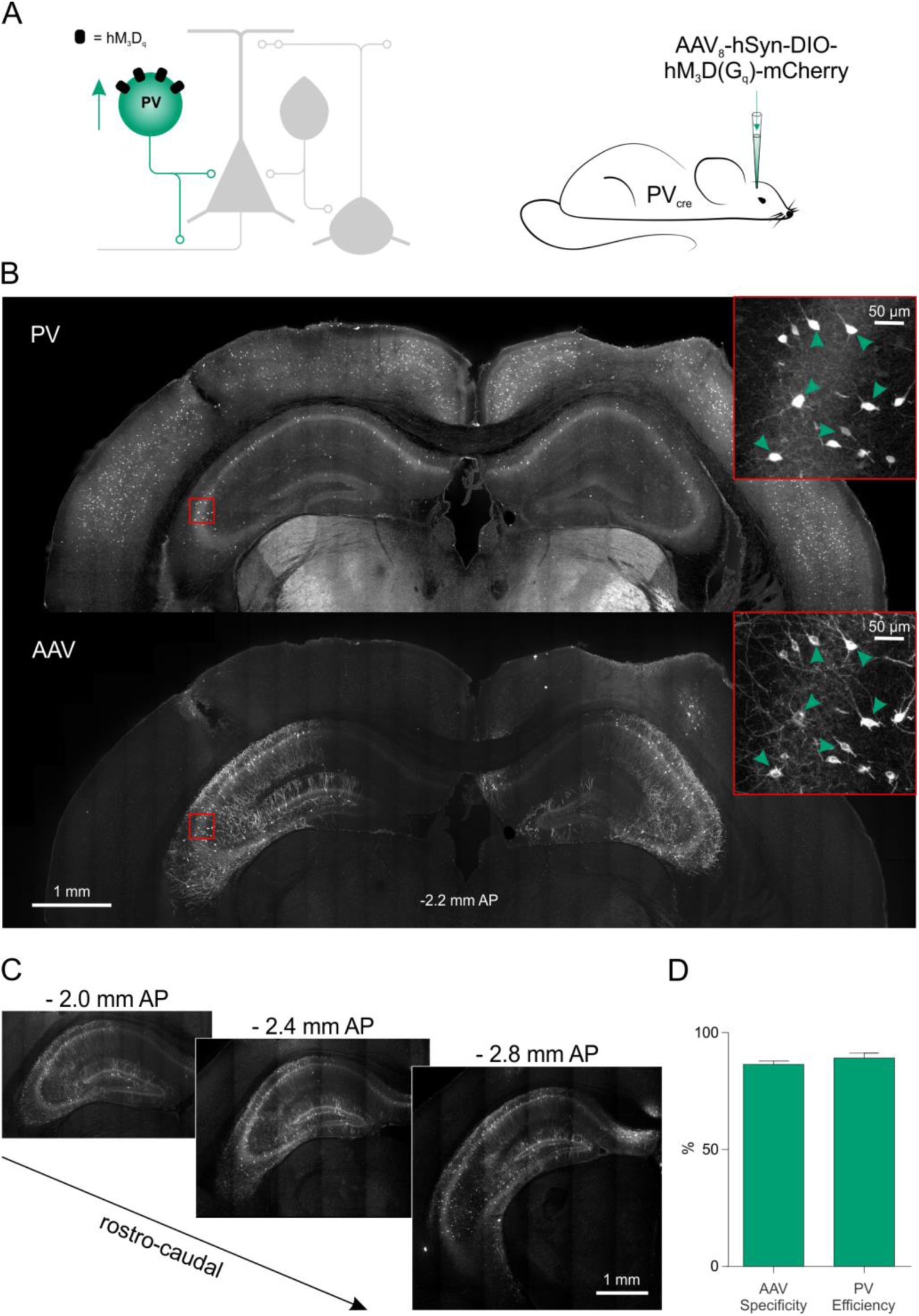
Excitatory DREADDs can be specifically and efficiently targeted to hippocampal PV interneurons in vivo. ***A***, To deliver hM_3_D_q_ receptors to PV interneurons in vivo (left), PV-cre mice received bilaterally injections of AAV8-hSyn-DIO-hM_3_D_q_-mCherry into the ventral and dorsal hippocampus (right). ***B***, Immunohistochemical characterisation confirmed that hippocampal PV interneurons (‘PV’, top) were efficiently transduced with hM_3_D_q_-mCherry (‘AAV’, bottom). Insets in red squares indicate cells co-expressing PV and hM_3_D_q_-mCherry (arrow heads). ***C***, Serial sections illustrate extensive AAV spread and hM_3_D_q_-mCherry expression throughout the hippocampus. AP, anteroposterior. ***D***, Population data showing high targeting specificity and efficiency of the hM_3_D_q_-mCherry in PV interneurons (N=7 animals).

To investigate the effect of activating PV interneurons on epileptiform activity in vivo, acute seizures were triggered by intra-hippocampal infusion of 4-AP. From 2-months after viral-mediated delivery of the hM_3_D_q_ receptor, the PV-cre mice were implanted with an intra-hippocampal infusion cannula. Animals underwent four seizure experiments, each separated by three days. Animals were randomised to receive an i.p. injection of either CNO or vehicle in their first experiment, and then alternated between CNO and vehicle for subsequent experiments. Each i.p. injection was delivered 15 minutes before the animal was placed in an arena and their freely moving behaviour was monitored for a period of 80 minutes using high-speed, high-definition cameras (**Fig. 7***A*). Following a 20-minute baseline period, 4-AP was infused directly into the hippocampus according to a spaced delivery protocol (three 4-AP infusions, each separated by 12 minutes; see Materials and Methods).

**Fig. 7.**
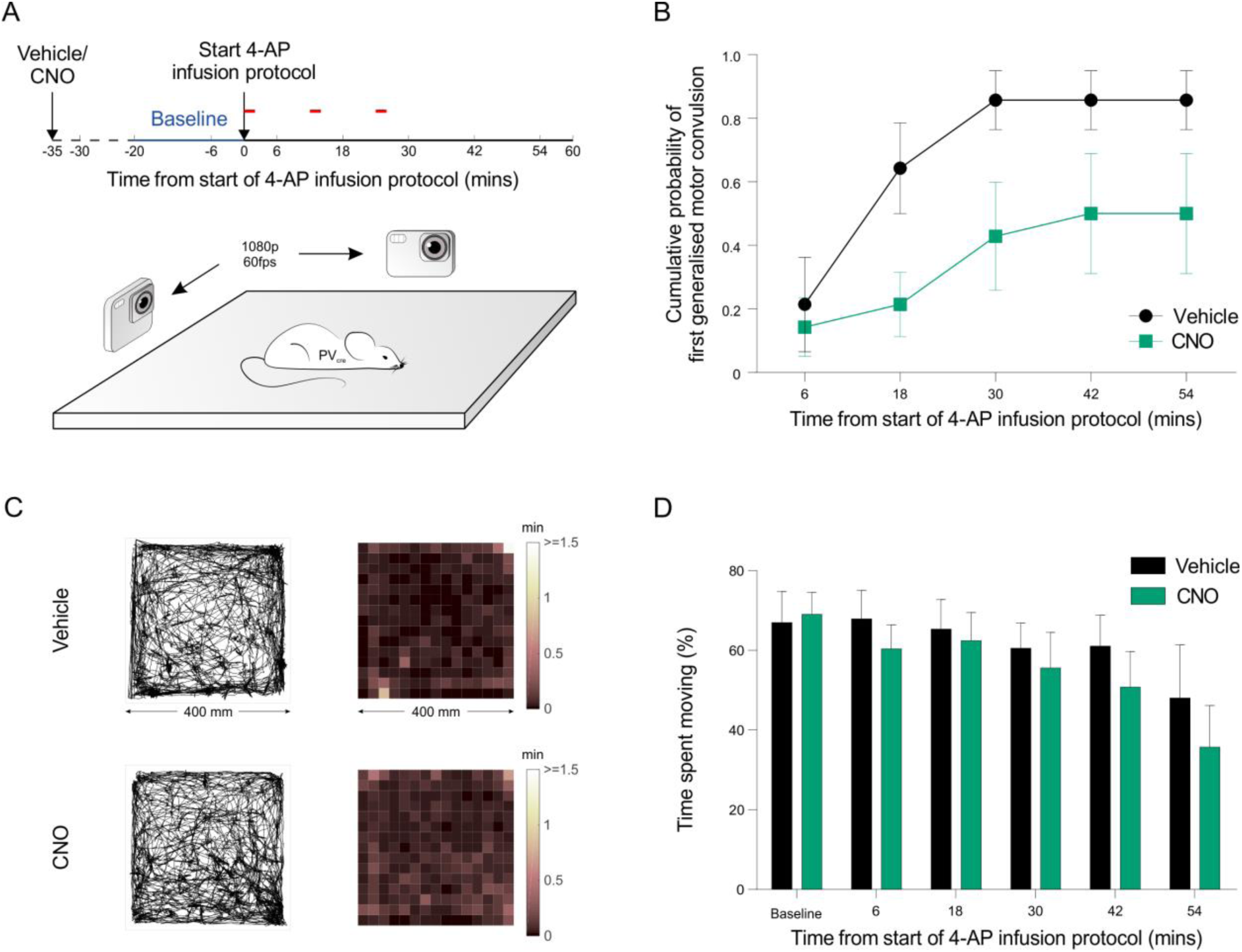
Chemicogenetic recruitment of hippocampal PV interneurons prevents generalised seizures in vivo. ***A***, Cartoon showing experimental design for assessing epileptiform activity in animals expressing the hM_3_D_q_ receptor in hippocampal PV interneurons. Fifteen minutes before behavioural monitoring, each animal received an i.p. injection of either vehicle control or CNO. Baseline behaviour was then monitored for 2o minutes (blue period), after which the intra-hippocampal 4-AP infusion protocol was started and monitoring continued for a period of 60 mins (red bars indicate the timing of each infusion). Behaviour was recorded by two high-speed, high-definition cameras oriented at right angles to one another, while a third camera tracked the animal from above. ***B***, The probability of reaching the first generalised motor convulsion (Racine 5 stage) was significantly reduced when CNO was administered compared to vehicle control (7 animals, non-cumulative data, t_(60)_=2.66, p=0.01, two-tailed t-Test corrected for multiple comparisons using the Holm-Sidak method). ***C***, For mice expressing the hM_3_D_q_ receptor in hippocampal PV interneurons, plots illustrate tracking data and corresponding spatial distribution of time spent across the behavioural arena (400 mm x 400 mm). Representative data are shown for an animal receiving an i.p. injection of vehicle (top) or CNO (bottom). In each case, data is shown for a 20-minute period following the start of the 4-AP infusion protocol. ***D***, There was no difference between the vehicle and CNO groups in terms of the percentage of time spent moving (N=10 vehicle and 11 CNO experiments, treatment: F_(1,104)_=1.686, p=0.197, two-way ANOVA).

For our initial analysis, we scored the time of the first generalised motor convulsion episode as the ‘generalised seizure threshold’ (i.e. reaching level 5 on the Racine scoring, at which point 4-AP infusions were terminated). The probability of reaching the generalised seizure threshold in the control experiments (i.e. vehicle injections) was highest between the second and third 4-AP infusions, with a probability of 0.43 ± 0.13 (N=12 experiments in 7 animals, reaching seizure threshold 12-24 minutes after the start of the 4-AP infusion protocol). In contrast, this probability was dramatically reduced to 0.07 ± 0.07 in animals that had received CNO injections to enhance the activity of their hippocampal PV interneurons (N=14 experiments in 7 animals). These effects were evident across the monitoring period, as shown by cumulative probability plots for generalised seizure threshold (**Fig. 7***B*). To assess whether this reduction in generalized seizures was associated with non-specific effects upon behaviour, we monitored the animals’ locomotor activity throughout the experiment via a camera mounted above the behavioural arena. These data revealed that the distribution of time spent throughout the arena was indistinguishable between the vehicle and CNO-treated groups (**Fig. 7***C,D*), supporting the conclusion that the CNO-mediated reduction in generalised seizures was not associated with a non-specific effect upon locomotor activity.

To provide a more detailed description of the epileptic seizure activity, each animal’s behaviour was scored blindly using the five-point Racine scale, at a sampling frequency of 1 Hz across at least 70 minutes per experiment (**Fig. 8***A*; see Materials and Methods). Racine categorization of epilepsy-related behaviour included orofacial clonic activity (Racine 1), head nodding (Racine 2), limb clonic activity (Racine 3), retreating/rearing with orofacial clonic activity (Racine 4) and rearing and falling and/or jumping (Racine 5). These analyses revealed that CNO-mediated recruitment of hippocampal PV interneurons caused a reduction in the frequency of epileptic behaviours across the Racine categories (**Fig. 8***B*). To characterise the temporal aspects of these effects, the Racine scoring scale was used to generate an integrated measure of epileptic-related behaviour that could be tracked over time (see Materials and Methods). This integrated measure revealed that CNO-mediated recruitment of hippocampal PV interneurons reduced the occurrence of all epileptic behaviours by 85% compared to controls (**Fig. 8***C*). In summary therefore, these results are consistent with the conclusion that chemicogenetic recruitment of PV interneurons is able to effectively supress seizure activity in vivo.

**Fig. 8.**
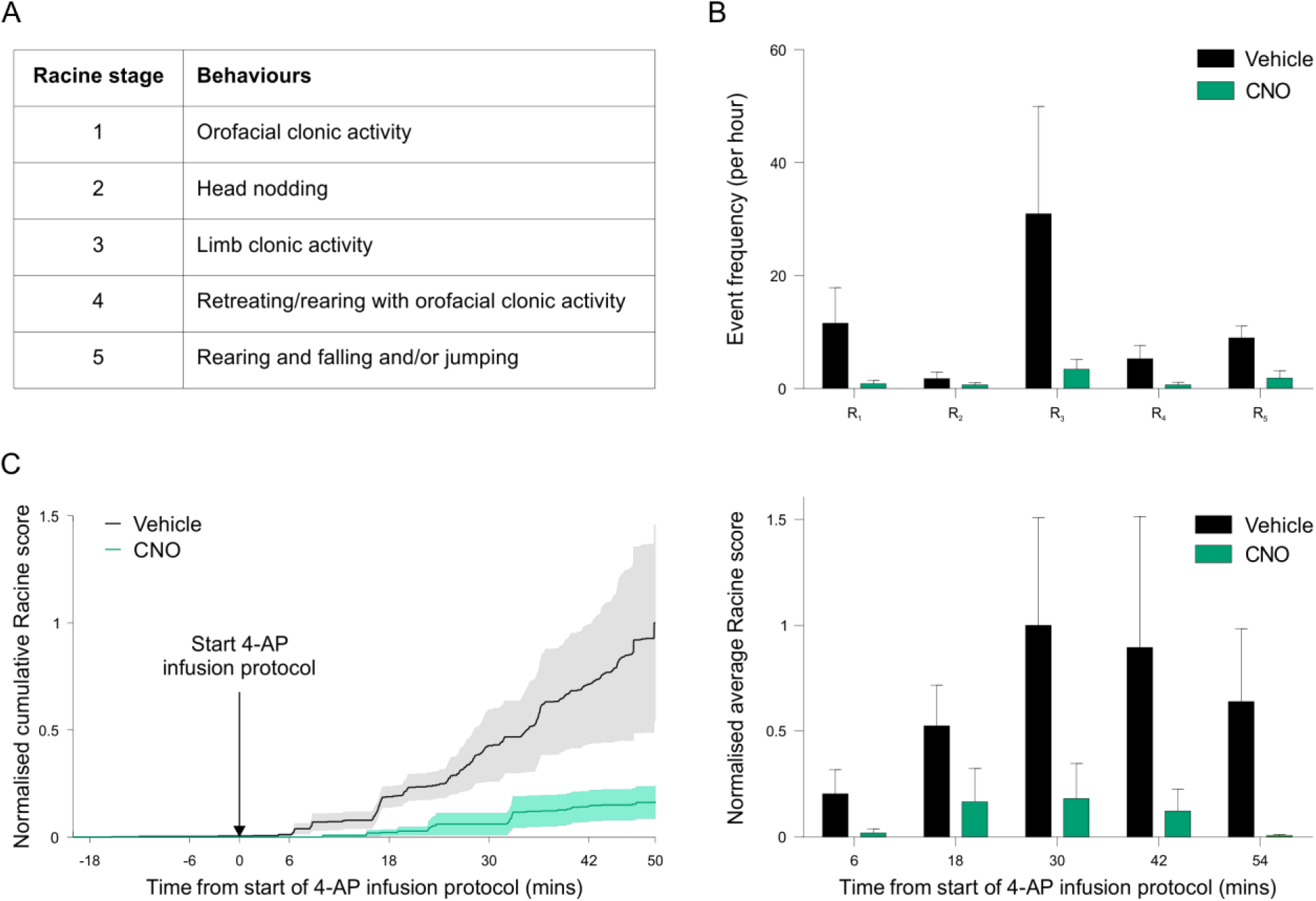
Chemicogenetic enhancement of hippocampal PV interneurons supresses epileptic behaviours in vivo. ***A***, >Table describing the Racine scale used to score the animal’s epileptic behaviour from video analysis offline. ***B***, Frequency of epilepsy-related behavioural events was significantly reduced when CNO was administered compared to vehicle across all Racine (R1-5) behaviours (N=12 vehicle and 10 CNO experiments, treatment: F_(1,100)_=5.161, p=0.0252, two-way ANOVA). ***C***, Epileptic behaviour, plotted as a cumulative score (left) or a non-cumulative score (right), was significantly lower following administration of CNO compared to vehicle (N=12 vehicle and 10 CNO experiments, treatment: F_(1,99)_=7.418, p=0.0076, two-way ANOVA).

## Discussion

Here we demonstrate the potential to generate anti-epileptic effects through the chemicogenetic enhancement of distinct populations of GABAergic interneurons. Using cre-recombinase mouse technology and viral delivery of excitatory DREADDs (Alexander et al., 2009; Roth, 2016; Smith et al., 2016), we demonstrate the efficacy of chemicogenetic recruitment of the PV, SST and VIP interneuron populations. Targeted whole cell recordings from both the pre-synaptic interneurons and post-synaptic principal neurons revealed that all three interneuron types increased their firing rate and synaptic output following CNO-mediated activation of hM_3_D_q_ DREADD receptors. Using the same in vitro system, we show that the interneuron populations exert different effects upon epileptiform activity. Chemicogenetic enhancement of either the PV or SST interneurons decreased spontaneous epileptiform activity, through a reduction in the frequency of SLEs. In contrast, enhancing VIP interneuron activity did not reduce total epileptiform activity. When accounting for the relative density of cells, PV interneurons generated the strongest anti-epileptic effect per cell. Finally, to confirm the potential of such an intervention strategy, chemicogenetic activation of PV interneurons was assessed in an in vivo model of temporal lobe seizures. Chemicogenetically boosting PV interneuron activity significantly lowered the level of epilepsy-related behaviours and the probability of reaching the threshold for generalised convulsions.

Previous work has shown that targeting pyramidal neurons with inhibitory DREADDs can mitigate seizure activity (Kätzel et al., 2014; Avaliani et al., 2016). The current study extends this by demonstrating that, although they make up a relatively small proportion of the network, chemicogenetically recruiting GABAergic populations can mediate robust anti-epileptic effects. Enhancing such endogenous inhibitory mechanisms may represent an attractive intervention strategy, as interneurons can exert widespread effects on the tissue and inhibitory circuits are recruited as excitatory network activity intensifies (Trevelyan et al., 2006; Derchansky et al., 2008; Schevon et al., 2012; Cammarota et al., 2013). The inhibitory network is therefore well-placed to respond to, and prevent, the spread of excitation from one or more sites in the tissue. However, given their diversity, the particular interneuron population that is targeted is likely to be important. In line with this prediction, our data show that chemicogenetic enhancement of VIP interneurons did not reduce overall seizure activity and in fact increased the duration of individual SLEs. This is in-line with observations that the preferential post-synaptic targets of VIP interneurons are other GABAergic interneurons (Acsády et al., 1996; Dávid et al., 2007; Pfeffer et al., 2013), and that activating VIP interneurons could prevent downstream GABAergic interneurons from counteracting seizure-related excitation. Consistent with such a disinhibitory effect, a recent study has shown that blocking VIP interneuron spiking can attenuate seizure-related activity (Khoshkhoo et al., 2017). More generally, this supports the conclusion that recruiting all inhibitory interneurons could generate complex effects and the importance of regulating specific interneuron types.

In contrast to VIP interneurons, PV and SST interneurons have been shown to preferentially target principal neurons (Klausberger et al., 2003). PV interneurons comprise mainly basket cells and axo-axonic cells, which synapse on the soma and axon-initial segment of pyramidal neurons (Klausberger and Somogyi, 2008). The dense axonal branching of a PV interneuron is therefore restricted to the pyramidal layer, but is broadly distributed with a transverse extent of ~1 mm in the rodent hippocampus. This can generate widespread synaptic inhibition, with each cell targeting the perisomatic region of around 1500-2000 pyramidal neurons (Freund and Buzsáki, 1996). SST interneurons meanwhile, mainly synapse on the dendrites of pyramidal neurons (Klausberger and Somogyi, 2008), where they regulate dendritic activation (Miles et al., 1996) and may account for as much as half of the firing rate increase following complete removal of inhibition (Lovett-Barron et al., 2012).

In our experiments, DREADD activation of either PV or SST interneurons resulted in pronounced increases in spike-evoked, post-synaptic inhibitory currents in pyramidal neurons and strong attenuation of spontaneous SLEs. For both cell types, we observed a ~50% reduction in the total SLE activity, which was driven by a decrease in the probability of SLE initiation. At the population level, enhancing SST interneurons generated the largest post-synaptic inhibitory currents in pyramidal neurons. Although when adjusted for cell numbers, we estimated that individual PV interneurons elicited equivalent post-synaptic inhibitory currents and a five-fold greater attenuation of SLE activity. These observations are consistent with evidence that GABAergic inputs to the axo-somatic region can exert particularly powerful inhibitory effects (Vu and Krasne, 1992; Cobb et al., 1995) and that PV interneurons are primarily responsible for mediating the synaptic restraint that can oppose the initiation and propagation of seizure activity (Cammarota et al., 2013; Paz and Huguenard, 2015). In the context of epilepsy, the targeting of PV interneurons may also be more preferable because there are reports that the SST interneuron population become depleted (Robbins et al., 1991; Cossart et al., 2001).

In agreement with these ideas and our in vitro results, we demonstrate that excitatory DREADDs in PV interneurons can generate potent anticonvulsant effects in vivo. Using a behavioural model in which epileptic seizures are initiated in the hippocampus, we show that chemicogenetic enhancement of PV interneurons within the hippocampus can significantly reduce the level of epilepsy-associated behaviours and the probability that the network crosses a threshold for generalised seizures. Interestingly, this reduction in epileptic behaviour was not associated with general effects on the animal’s locomotor activity, suggesting that the chemicogenetic enhancement of PV interneurons becomes evident when the network enters a pre-epileptic state, rather than causing widespread effects upon network activity. More generally, the data illustrate that chemicogenetically enhancing a specific interneuron population can produce effective anticonvulsant effects.

A series of studies using rodent models have successfully applied optogenetic approaches for disrupting and dissecting epileptiform networks. Like chemicogenetic approaches, optogenetics lends itself to cell-type targeting and this work has helped define the role of different interneuron types. For example, it has been shown that when combined with real-time seizure detection methods, temporally controlled optogenetic activation of interneuron populations can provide effective disruption of epileptic activity (Krook-Magnuson et al., 2013, 2014; Ledri et al., 2014; Sessolo et al., 2015; Assaf and Schiller, 2016; Khoshkhoo et al., 2017). This work has also revealed that optogenetic activation of interneurons can actually initiate epileptiform activity, in a manner that may depend on the network state (Sessolo et al., 2015; Shiri et al., 2015; Yekhlef et al., 2015; Assaf and Schiller, 2016; Chang et al., 2018). This phenomenon seems to be associated with the pulsed light-activation and enforced synchronisation of interneuron activity, which can then synchronise the network by inducing time-locked post-inhibitory rebound spiking (Jefferys et al., 2012; Sessolo et al., 2015; Chang et al., 2018). In addition to these temporal aspects, optogenetic strategies face other challenges for disrupting seizures, including the delivery of light to structures that may be deep within the brain, or to cells that may be distributed over large regions.

Chemicogenetic intervention strategies may mitigate these issues, such as the potential to modulate cellular activity on larger spatial and temporal scales. Effects from chemicogenetics can be coordinated across large areas of tissue, due to the systemic delivery of the activating drug (Roth, 2016). Furthermore, the fact that DREADDs are G-protein coupled receptors and act through endogenous cellular mechanisms, may avoid unwanted effects associated with artificial synchronisation of the network (Wang et al., 2017). However, their reliance upon the cell’s own intracellular machinery may limit the potential for chemicogenetics to modulate the activity of specific cell types. Whereas optogenetic strategies have the power to use light energy to move ions against concentration gradients, chemicogenetic strategies must rely upon endogenous mechanisms, at least in their current form. Also, there remain questions about how inert the designed drugs are for the selective activation of the DREADDs, and whether these need to be refined further (Gomez et al., 2017). Any cell-targeted strategy must also consider how the contributions of particular cell types may change in epilepsy. Cell types may be lost or change their signalling at different stages of the disease (Robbins et al., 1991; Cohen et al., 2002; Wang et al., 2017), their contribution may depend on their location relative to the epileptic focus (Sessolo et al., 2015) or their position in the epileptic circuit (Paz and Huguenard, 2015; Bui et al., 2018), and their effects may change dynamically during an individual seizure (Ellender et al., 2014). For these reasons, multiple strategies may be required, perhaps using one strategy for pathologically affected cells within the epileptic focus and another strategy for surrounding healthier circuits.

In conclusion, the current work supports the use of selective chemicogenetic targeting of the inhibitory system as an approach to disrupt seizure activity. Such a cell-specific pharmacological strategy has the attraction of being controllable and yet avoiding the system-wide effects of drugs that enhance GABA-mediated inhibition in a non-cell selective fashion. Future work will be required before such cell-targeted strategies can be adopted in a translational context, but increasing support is being provided for this general principle.

## Acknowledgements

The research leading to these results has received funding from the European Research Council under the European Community’s Seventh Framework Programme FP7/2007-2013, ERC Grant Agreement 617670. A.C. was supported by a Wellcome Trust Doctoral Fellowship [102364/Z/13/Z]. A.S.I. was supported by a Junior Research Fellowship in Medical Sciences from University College Oxford. We thank Miruna Rașcu for help in processing the mouse tracking data. We would also like to thank Professor Peter Somogyi for providing advice and the antibody against vasoactive intestinal polypeptide.

**Author contributions**
A.C., A.S.I. and C.J.A. conceived the project. All authors contributed to the design of the experiments. A.C. and M.S. performed the experiments. A.C. analysed the data and prepared all figures. A.C. A.S.I. and C.J.A. wrote the manuscript. All authors reviewed the manuscript.

